# Transcriptome-wide association supplements genome-wide association in *Zea mays*

**DOI:** 10.1101/363242

**Authors:** Karl A. G. Kremling, Christine H. Diepenbrock, Michael A. Gore, Edward S. Buckler, Nonoy B. Bandillo

## Abstract

Modern improvement of complex traits in agricultural species relies on successful associations of heritable molecular variation with observable phenotypes. Historically, this pursuit has primarily been based on easily measurable genetic markers. The recent advent of new technologies allows assaying and quantifying biological intermediates (hereafter endophenotypes) which are now readily measurable at a large scale across diverse individuals. The potential of using endophenotypes for dissecting traits of interest remains underexplored in plants. The work presented here illustrated the utility of a large-scale (299 genotype and 7 tissue) gene expression resource to dissect traits across multiple levels of biological organization. Using single-tissue- and multi-tissue-based transcriptome-wide association studies (TWAS), we revealed that about half of the functional variation for agronomic and seed quality (carotenoid, tocochromanol) traits is regulatory. Comparing the efficacy of TWAS with genome-wide association studies (GWAS) and an ensemble approach that combines both GWAS and TWAS, we demonstrated that results of TWAS in combination with GWAS increase the power to detect known genes and aid in prioritizing likely causal genes. Using a variance partitioning approach in the independent maize Nested Association Mapping (NAM) population, we also showed that the most strongly associated genes identified by combining GWAS and TWAS explain more heritable variance for a majority of traits, beating the heritability captured by the random genes and the genes identified by GWAS or TWAS alone. This improves not only the ability to link genes to phenotypes, but also highlights the phenotypic consequences of regulatory variation in plants.

**Author summary:** We examined the ability to associate variability in gene expression directly with terminal phenotypes of interest, as a supplement linking genotype to phenotype. We found that transcriptome-wide association studies (TWAS) are a useful accessory to genome-wide association studies (GWAS). In a combined test with GWAS results, TWAS improves the capacity to re-detect genes known to underlie quantitative trait loci for kernel and agronomic phenotypes. This improves not only the capacity to link genes to phenotypes, but also illustrates the widespread importance of regulation for phenotype.

## Introduction

Discovery of variation that underlies quantitative traits remains central to the genetic improvement of agricultural species. Functional variation can alter coding sequence or act to regulate an intermediate phenotype. Regulating the abundance of phenotypic intermediates, like mRNA expression or protein level, provides a more spatially and temporally subtle target for selection than coding sequence changes, which are more likely to be pleiotropic and therefore maladaptive [1]. Thus, regulatory variation is the frequent target of both natural and artificial selection that shapes genomes across life, including domesticated plants [1-3]. It is likely that about half of functional variation is regulatory [4-7]. It should also be noted that regulation can take place at any biological level of organization from the epigenetic state [8], to gene expression [4,9,10], to ribosome occupancy [11], to metabolites [12], to protein abundance [13,14], furnishing multiple levels at which intermediate and terminal phenotypes can be associated.

In standard genetic mapping approaches, like association or linkage mapping, associations between genetic markers and terminal phenotypes of interest are tested for significance (black arrow, Fig 1). However, multiple levels of biological organization exist between the DNA sequence and the terminal observed phenotypic outcomes, enabling trait dissection to be done between intermediate levels of biological organization (hereafter endophenotypes, designated by an orange and red arrow in Fig. 1). Associating endophenotypes with terminal phenotypes predates the use of molecular genetic markers for mapping. The use of linked observable traits and isozyme migration patterns are examples of tying markers from biological intermediates to terminal phenotypes of interest. Similarly, just as relationships between individuals can be calculated from molecular genetic markers [15], endophenotypic similarity from isozyme markers can also be used to quantify relatedness [16]. These same principles have recently been extended to phenotypic prediction guided by metabolites [12] or by expression dysregulation [17]. However, the use of molecular intermediates, which are now readily measurable at large scale across diverse individuals, remains underexplored in plants for the inverse task of causal inference.

**Figure 1.**
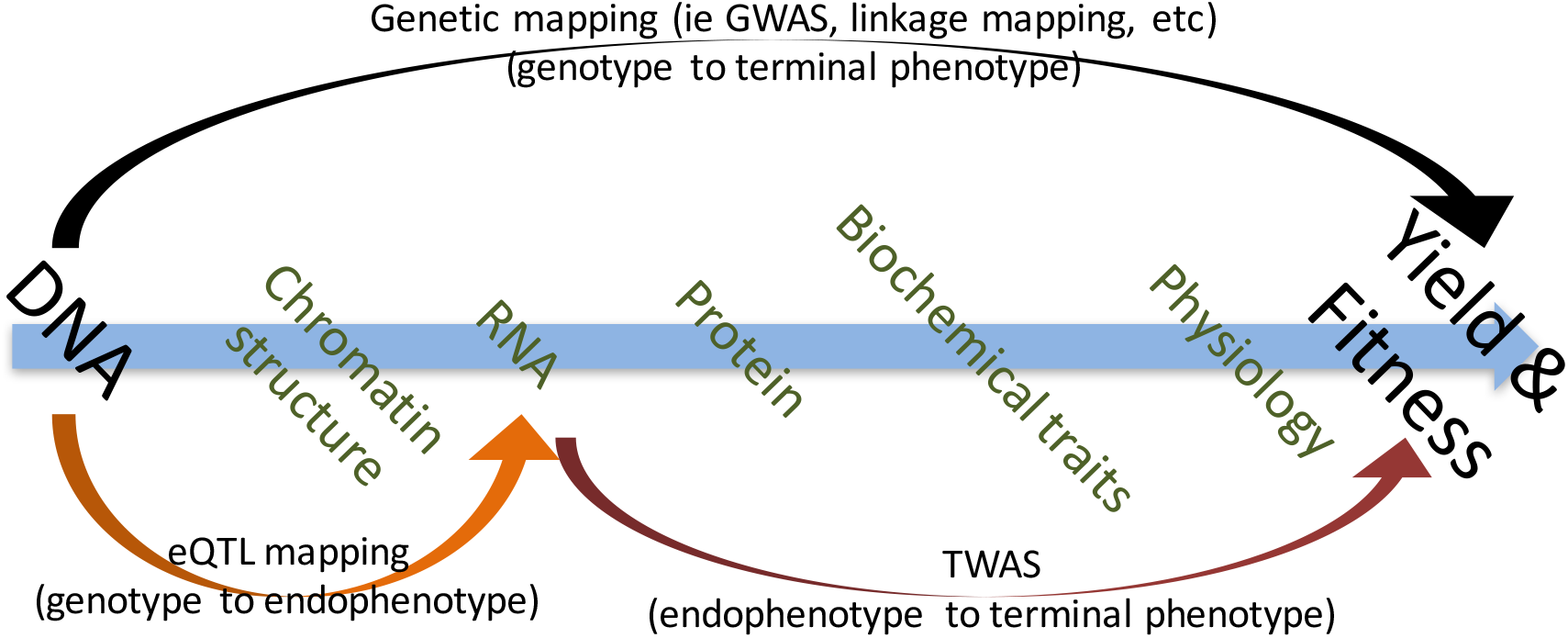
Levels of biological organization between the ultimate cause of genetics and the terminal phenotypic outcomes can be exploited individually to improve power and inference of biological mechanism. Genotype can be linked to endophenotype as in eQTL or protein QTL (pQTL), or endophenotype can be linked to terminal phenotype by methods like TWAS.

Associating endophenotypes with terminal phenotypes has multiple distinct advantages. First, while genetic mapping is dominated by the covariance structure of neighboring SNPs and complex haplotypes, endophenotype provides orthogonal information that often permits inference about biological mechanism, which may not be possible from genetic variants alone. Second, genetic mapping often points to intergenic [18] regulatory variants that are not within the coding sequence of the gene that alters the phenotype [4]. Therefore, an association signal cannot directly be tied to a corresponding gene and may even be in the body of a second unrelated gene [19] or in the case of synthetic association, between multiple true causal variants affecting different genes. Association tests with intermediate expression phenotypes do not suffer from these limitations. Third, the abundance of endophenotypes is largely independent of linkage disequilibrium (LD), unlike in the case of genetic markers. In other words, even multiple genes that are perfectly linked, and thus not independently observable in separate individuals, can be prioritized for association with a trait because their expression patterns are independent. This is of greatest utility in species where linkage disequilibrium is extensive or where making high-resolution mapping populations is not feasible.

Intermediate phenotypes, like expression, can also integrate the signal from changes in multiple components of a network, which may not be individually detectable either because their effects are small or changes to the peripheral network components occur at low frequencies. Similarly, intermediate phenotypes can integrate a phenotypic signal from underlying genetic variants for which low frequencies preclude direct detection. The most deleterious of variants are expected to segregate at the lowest frequencies [20,21] and, thus, escape detection by mapping without prohibitively large sample sizes. However, rare deleterious variants can be expected to drive common maladaptive patterns in intermediate phenotypes that are thus more easily detected through endophenotype association tests like transcriptome-wide association studies (TWAS) [22,23]. It is precisely the shared effect of different alleles on intermediate phenotypes like expression that is exploited in expression-based prognoses for cancers [24,25] and which TWAS can also leverage.

Here, we illustrate the power of using gene expression endophenotypes measured in a large 299-individual, seven-tissue gene expression resource [17] collected from the Goodman maize diversity panel [15]. Expression levels are correlated with terminal phenotypes in TWAS [22,23] and then combined with genotype-based associations from GWAS. The method is proven here in a maize inbred diversity panel [15], which has been widely used to dissect the architecture of dozens of traits of varying complexity [26-29], and is now made more powerful to detect loci at the gene level through the integration of transcriptome-wide with genome-wide associations.

Related work in maize that relies on associating expression differences directly with phenotype using a Bayesian method, called expression read depth GWAS (eRD-GWAS), has been published recently [30]. This work used 369 maize samples from which shoot apex RNA was collected. Beyond the difference in frequentist vs Bayesian approaches, our study also exploits expression measurements from seven tissues in a multiple-regression-based TWAS and integrates the signal from TWAS and GWAS into a more powerful combined test which can be readily visualized as a Manhattan plot. We also compare the power of each model based on the ability to detect known genes, and the capacity to explain variance in a separate population, which differs from the approach of the previous study [30]. To make this comparison we use the maize NAM population [31], which has the advantage of being largely independent of the diversity panel [15] in which detection was performed.

We assess the efficacy of TWAS by quantifying the capacity to identify previously identified genes, and by the fraction of phenotypic variance explained [5,6] by the most strongly associated genes, and compared the TWAS results with GWAS and an ensemble approach combining both TWAS and GWAS. We illustrate that the results of TWAS are a valuable supplement to GWAS mapping that aids in prioritizing likely causal genes when both methods are used in a combined test.

## Results

To test the utility of expression data in dissecting quantitative traits in maize, we performed single-tissue-based and multi-tissue-based TWAS [22] and compared these results with GWAS results, and an ensemble approach combining GWAS and TWAS results using the Fisher’s combined test. In TWAS, expression levels across seven tissues from a maize diversity panel [15] were used individually and together in a multiple regression as independent variables and correlated with previously measured phenotypes for maize kernel traits, including 30 grain carotenoid abundance traits [28], 20 tocochromanol abundance traits [27], and 22 field-measured agronomic traits [32].

### Integrating TWAS with GWAS improves power for identifying and prioritizing known genes

To assess the relative power of each method to detect known genes, we counted the number of known genes identified in the top 1% ranked genes (based on p-values) found by each method for each trait. This identification of known genes among the top 1% of hits for each method measures how often known genes appear in the tail of the distribution of detected genes and avoids direct comparisons of p-values between differently powered and structured tests that rely on continuous (TWAS) or discrete (GWAS) independent variables.

As shown in Tables 1, 2, S1, S2, the combined test performs better than either genotype-based or expression-based tests alone for both classes of traits, with 30 total detections of known genes among the top 1% of associations across tocochromanol and 75 detections of putative carotenoid related genes [28] when using the carotenoid traits. Using the tocochromanol and carotenoid lists from [25, 36] genes are detected more often in each of the tocochromanol and carotenoid trait classes when using the combined method. However, the Fisher’s combined test of GWAS results with the multi-tissue TWAS results did not perform better. The detection rate was consistently higher for kernel-based TWAS over the multi-tissue TWAS, most likely because the tocochromanol and carotenoid traits are predominantly controlled by gene expression in the kernel.

**Table 1.** Summary of total and unique known gene detections in top 1% of results across tocochromanol traits by kernel TWAS with PEERS and PCs, multi-tissue TWAS with PCs, MLM GWAS, Fisher’s combined test of kernel TWAS with PEERS and PCs and MLM GWAS, and Fisher’s combined test of multi-tissue TWAS with PCs and MLM GWAS. There are 14 previously known tocochromanol genes in maize [36]. On the left half of the table the number of detections exceeds the number of known genes because a gene is counted as detected each time it is in the top 1% of associations for the 20 tocochromanol component traits.

| Test | Detections of known genes in top 1% of hits<br>across tocochromanol traits |
| --- | --- |
| TWAS | 14 |
| multiTWAS | 13 |
| GWAS | 21 |
| FisherGWASTWAS | 30 |
| FisherGWASmultiTWAS | 27 |

**Table 2.** Summary of total and unique putative carotenoid gene [28] detections in top 1% of results across carotenoid traits by kernel TWAS with PEERS and PCs, multi-tissue TWAS with PCs, MLM GWAS, Fisher’s combined test of kernel TWAS with PEERS and PCs and MLM GWAS, and Fisher’s combined test of multi-tissue TWAS with PCs and MLM GWAS. On the left half of the table the number of detections exceeds the number of known genes because a gene is counted as detected each time it is in the top 1% of associations for the 30 carotenoid component traits.

| Test | Detections of candidate genes in top<br>1% of hits across carotenoid traits |
| --- | --- |
| TWAS | 38 |
| multiTWAS | 32 |
| GWAS | 55 |
| FisherGWASTWAS | 75 |
| FisherGWASmultiTWAS | 58 |

We also compared the methods at the level of single traits. To determine how the combined method prioritizes genes that are not detected in the individual TWAS and GWAS methods and aggregates genes that are detected by only one method, we plotted the results across models for each individual trait. In Fig. 2 we plotted the signals mapped for the zeaxanthin trait by (a) the mixed linear model (MLM) GWAS, (b) TWAS in kernels, (c) GWAS colored by TWAS significance, (d) the Fisher’s combined test of MLM GWAS and single tissue (kernel), and (e) Fisher’s combined test of MLM GWAS and multi-tissue TWAS results (*see* methods). Note that points representing SNPs from the MLM GWAS model in (c) and (a) are identically placed, but in (c) they are colored by TWAS significance. The top five genes detected by each method are labeled (a is not individually labeled because the points and top five genes are identical to those in plot c) and previously detected genes found by [28] are highlighted in red. As shown by the TWAS results plotted in Fig.2 b, the known expression-regulated gene *crtRB1* has expression which is most strongly correlated (r=0.309, p=2.84e-5) with zeaxanthin abundance in our TWAS model that includes genetic and expression-derived covariates [see methods]. *crtRB1* is not among the top MLM GWAS-detected genes in our study, but the detection of *crtRB1* by kernel TWAS is consistent with previous results [28,33], highlighting this gene’s role as a principle determinant of grain carotenoids which acts through variable expression.

**Figure 2.**
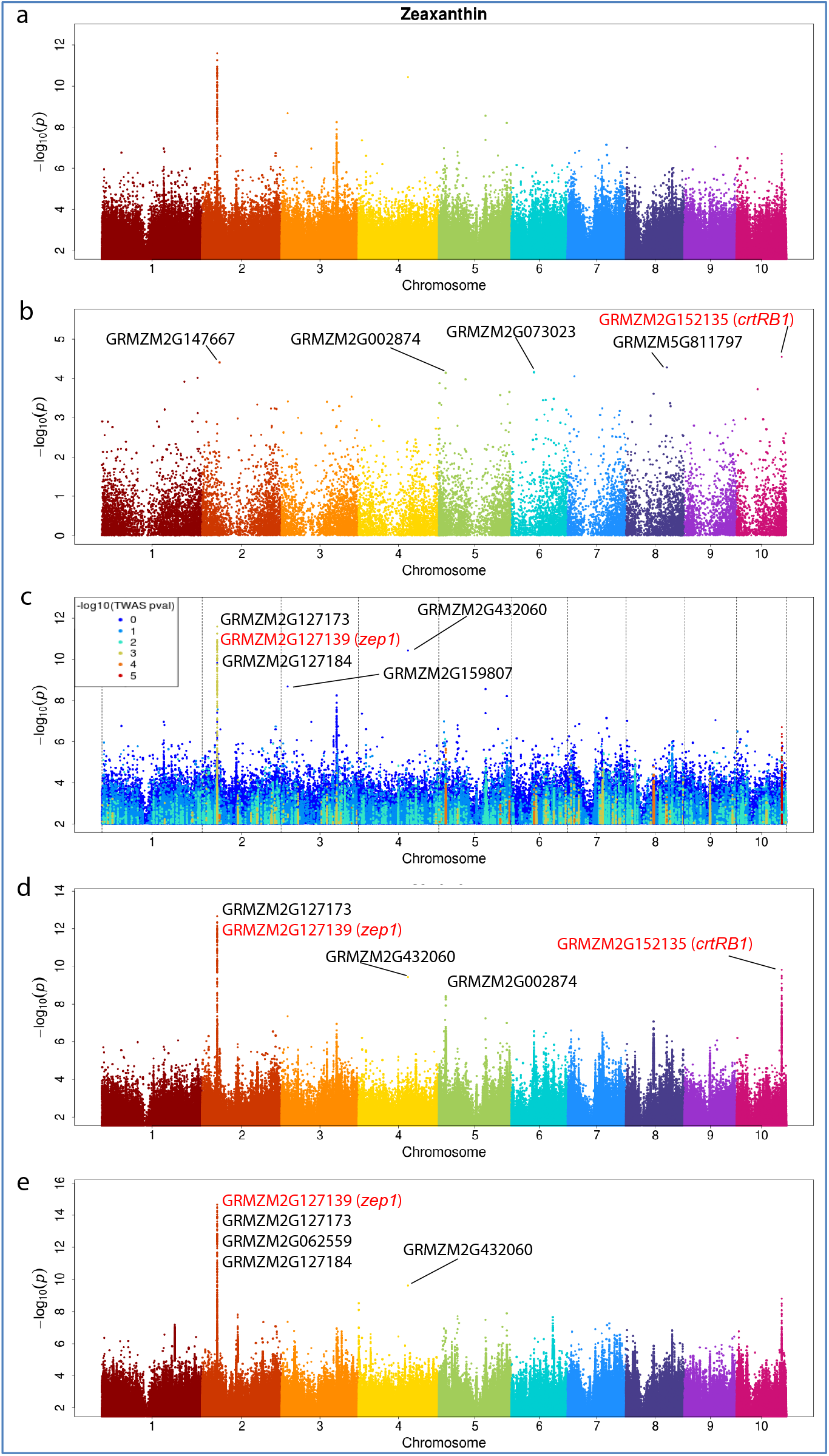
Manhattan plots of zeaxanthin abundance **a** mixed linear model GWAS accounting for kinship, **b** kernel TWAS with PEER and genetic MDS PC covariates, **c** MLM colored by TWAS significance, and **d** Fisher’s combined model of MLM and TWAS p-values using kernel expression. **e** Fisher’s combined model of MLM and multi-tissue TWAS p-values. The top five most associated genes are labeled and previously identified genes [28] are highlighted in red.

As is clear in Fig.2a,c another zeaxanthin-implicated gene, zeaxanthin epoxidase, *zep1*, is detected by GWAS in our study [28]. *zep1* expression is correlated (r= 0.232, p=0.0014) with zeaxanthin abundance, but it is not among the fifty most significantly associated genes in our TWAS results, and would not be prioritized by TWAS alone. However, within the peak covering *zep1* in Fig.2 a,c the markers most strongly associated with zeaxanthin from the MLM GWAS results prioritize a different gene first, GRMZM2G127123, which lacks a known function. The linkage-independent kernel TWAS results also show nearly equal support for both genes, providing evidence that GRMZM2G127123 (r= 0.218, p=0.0025) and *zep1* (r= 0.232, p=0.0014) both affect zeaxanthin abundance. Both Fisher’s combined models using the single-tissue and multi-tissue TWAS results also support the importance of both genes.

To test the capacity of the TWAS, GWAS and combined methods to re-identify genes known to underlie QTL for another trait class, we examined the detected genes for the total tocotrienol trait measured by [27]. In Fig. 3 the most strongly associated variant identified by GWAS is on chromosome 9 nearest a gene of unknown function, GRMZM2G431524. However, as is illustrated in the MLM GWAS Manhattan plot in which points are colored by TWAS significance (c), the other points in the chromosome 9 peak are near other genes known to underlie QTL whose expression is variably associated with total tocotrienol abundance. These second and third most strongly associated genes based on proximity to the most significant markers identified by GWAS are GRMZM2G345544 (function unknown) and *hggt1*, which has been previously tied to total tocotrienol content [27], and is essential for tocotrienol biosythesis. However, because *hggt1* expression is most strongly correlated with total tocotrienol measurements from among these first three genes in the chromosome 9 peak, the combined test using single tissue and multiple tissues of expression data prioritizes the known gene *hggt1* suggesting it is the functional gene in this region, consistent with previous evidence. This illustrates how the supplementary information from expression associations prioritizes likely causal genes that are not among the top hits of either individual expression or genotype-based methods.

**Figure 3.**
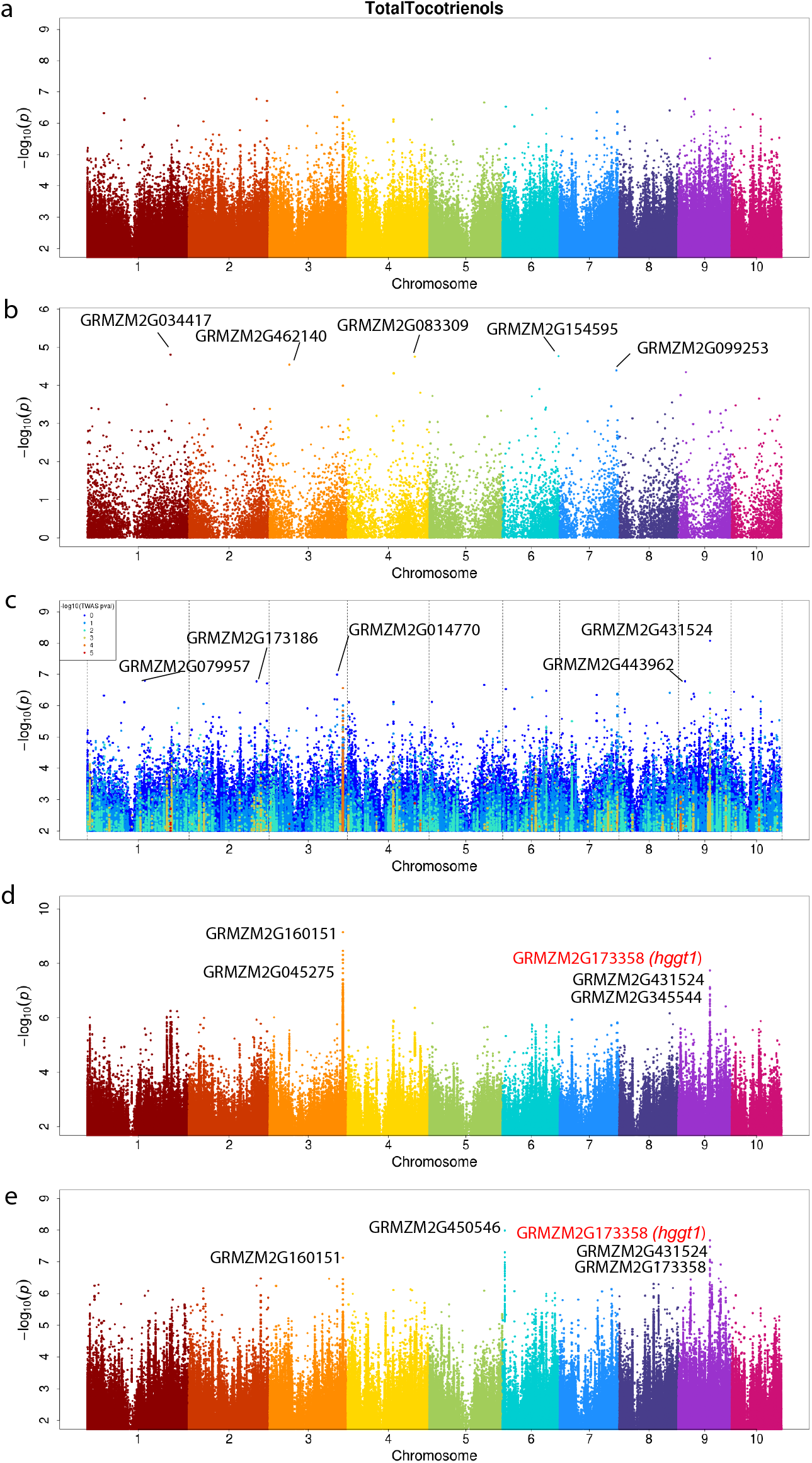
Manhattan plots of total tocotrienol abundance **a** mixed linear model GWAS accounting for kinship, **b** kernel TWAS with PEER and genetic MDS PC covariates, **c** MLM colored by TWAS significance, and **d** Fisher’s combined model of MLM and TWAS p-values using kernel expression. **e** Fisher’s combined model of MLM and multi-tissue TWAS p-values. The top five most associated genes are labeled and previously identified genes [36] are highlighted in red.

### Variance component estimation from TWAS- and GWAS-detected genes

To further assess the capacity of each method to correctly identify genes affecting each trait, an independent variance partitioning approach [5,6,35] was also performed. Using variants in a 1 Mb window around the ten top ranked genes identified in the Goodman diversity panel [15] by GWAS alone, TWAS alone and the combined method, separate kinship matrices were calculated. These relationship matrices were fit as random effects in separate models of phenotypic variance explained for traits measured in the NAM population, which is independent of the Goodman diversity panel in which the various mapping strategies were performed. The additive genetic variance explained by the variants underlying each kinship matrix was calculated providing an estimate of heritability explained by the genes identified by each method.

Using variance partitioning across all NAM families, we found some advantage for including expression data in detecting likely functional regions of the genome (Fig. 4). Among the tocochromanol kernel traits (Fig. 4a), eight out of ten traits exist in which TWAS or the Fisher’s combined method is superior to GWAS alone (Fig. 4a). Heritable variance explained on a per trait basis by either the TWAS or the Fisher’s showed about 25% improvement on average over the MLM GWAS, with notable advantage being for alpha-tocotrienol (40%), gamma-tocotrienol (41%) and total tocopherol (43%). For more complex field-based agronomic traits, the multi-tissue TWAS or Fisher’s combined method also showed advantage over GWAS alone in 16 out of 22 agronomic traits (Fig. 3b). On average, the multi-tissue TWAS had 24% improvement over GWAS alone while the FisherGWASmultiTWAS had notable advantage for kernel number (24%), leaf width (15%), and node number below ear (19%). Based on mean heritable variance across traits per trait class, the combined Fisher’s test explained the most heritability among the models; it showed 4-8% improvement for the tocochromanol kernel traits (Fig. 4a, the top right horizontal barplot). However, little improvement was observed for agronomic traits likely due to trait complexity (Fig. 4b, the top right horizontal barplot). Because previously known genes are more often re-identified in the top 1% of hits by combining GWAS and TWAS (Table 1), the variance explained by markers near detected genes also reflect this advantage on heritability with known oligogenic architecture.

**Figure 4.**
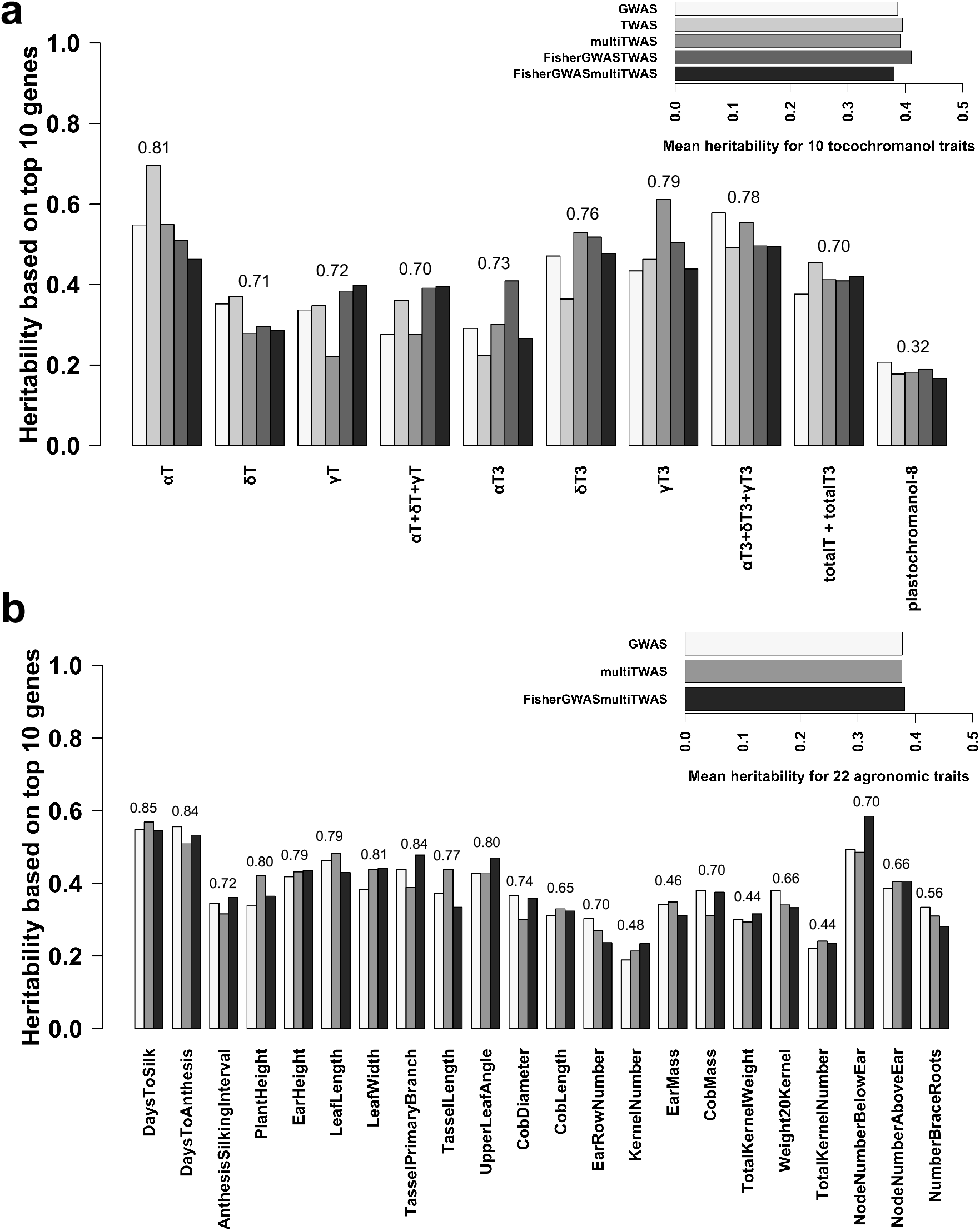
Variance partitioning of heritable variation using all NAM families. Vertical barplots represent the heritability estimated from kinship matrices made from the genetic regions adjacent to the top 10 ranked genes mapped by MLM GWAS, kernel-based TWAS, multi-tissue TWAS, the Fisher’s combined test of the MLM GWAS + kernel-based TWAS, and the Fisher’s combined test of the MLM GWAS + multi-tissue TWAS. Horizontal barplots compare model based on mean heritability across traits per trait class. Heritability explained by using all SNPs for each trait was put at the top of each grouped barplot.

We further tested the heritability explained by the top ten ranked genes identified by each method using family-based variance partitioning (Fig. 5). Heritable variance was decomposed for each NAM family, giving 24 independent tests of variance partitioning for each trait tallying a total of 3,840 independent tests (24 families * 5 models * 32 traits). To evaluate the best winning model for each trait, we took the sum of heritable variation across 24 NAM families (hereafter, summed heritability). Based on the same of set of genes identified from each model, our results illustrate the variability on heritability among families for both tocochromanol (Fig. S1, Fig. 5a) and agronomic traits (Fig. S2, and S3; Fig. 5b). For α-tocotrienol which is an oligogenic trait, the FisherGWASTWAS method explained the most heritability in 18 out of 24 NAM families (Figure S1a), giving a four-fold advantage on summed heritability over either GWAS or TWAS alone (Figure S1b; Fig. 5a). As is evident in the top right horizontal barplot (Fig. 5a), the FisherGWASTWAS method captured the most summed heritability in 10 tocochromanol traits, consistent with what we found in variance partitioning using all NAM families for tocochromanol traits (Fig. 4a). On a per trait basis, we note that the kernel-based TWAS or the FisherGWASTWAS was the winning method for eight out of 10 tocochromanol traits. We do see a similar pattern in 19 out of 22 field-based complex traits in which either the multi-tissue TWAS or FISHERGWASmultiTissueTWAS explained the most heritability (Fig. 5b). We see greater advantage of the FISHERGWASmultiTissueTWAS over the GWAS MLM for tassel primary branch (54%), cob length (103%), kernel number (112%), ear mass (98%) and total kernel weight (106%) (Fig. 5a). For the more complex traits such as plant height, the multi-tissue TWAS was the winning model, which explained about two-fold higher heritability than the GWAS alone (Fig. 5b, Fig. S3). We found that in 16 NAM families, the multi-tissue TWAS explained the most heritability among other models for plant height. Based on total summed heritability across 22 agronomic traits (top right horizontal barplot in Fig. 5b), the FISHERGWASmultiTissueTWAS and multi-tissue TWAS showed a 15% and 17% improvement in heritability explained over the GWAS MLM alone, respectively.

**Figure 5.**
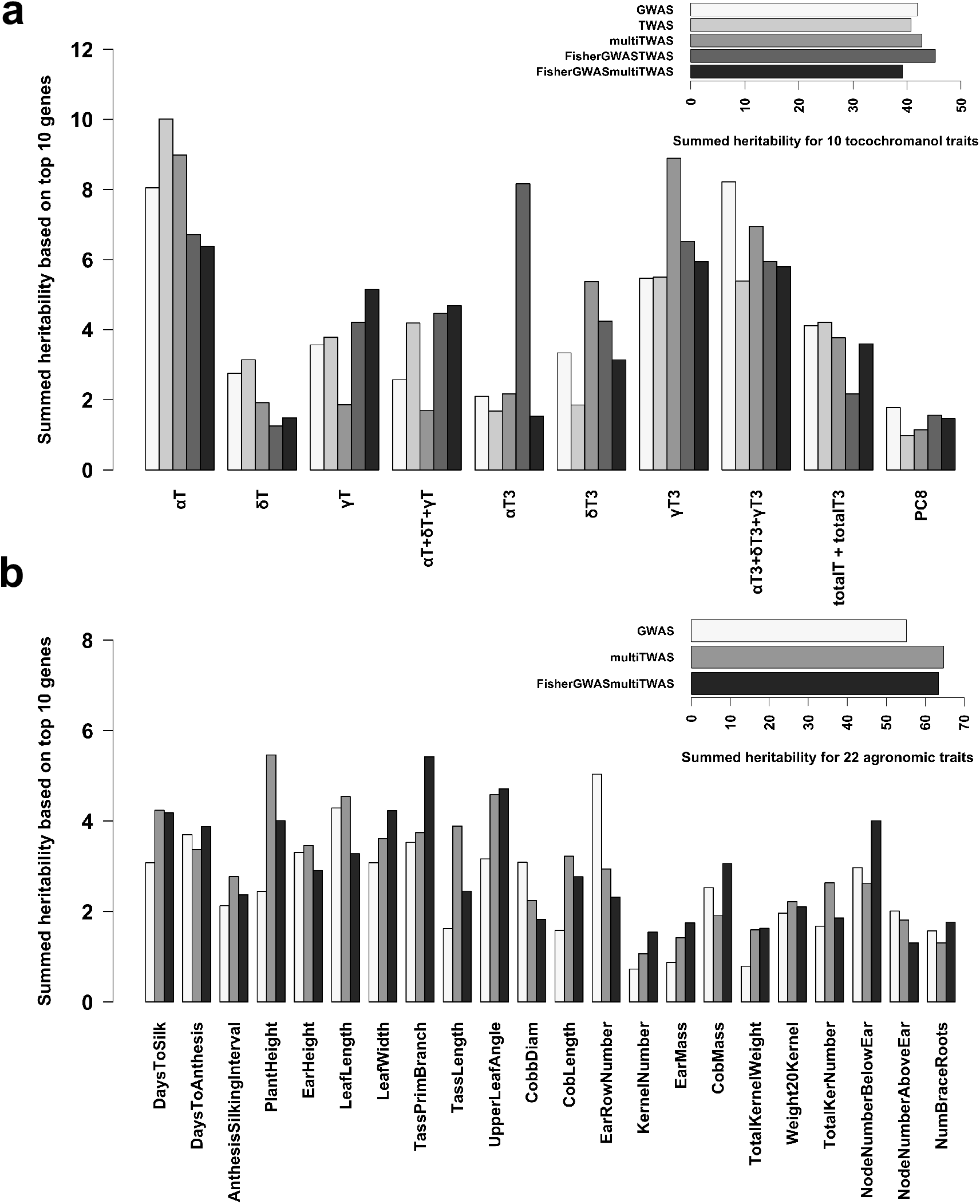
Family-based variance partitioning on individual NAM family. Heritability for each trait was estimated for each of 24 NAM families using kinship matrices made from the genetic regions adjacent to the top 10 ranked genes mapped by MLM GWAS, kernel-based TWAS, multi-tissue TWAS, the Fisher’s combined test of the MLM GWAS + kernel-based TWAS, and the Fisher’s combined test of the MLM GWAS + multi-tissue TWAS. There were a total of 24 independent tests for each trait-model combination. Heritability estimates were then added together (hereafter, summed heritability) for **a** tocochromanol traits and **b** agronomic traits. Horizontal barplots compare model based on total summed heritability across traits per trait class.

## Discussion

By far the majority of efforts to dissect the architecture of terminal phenotypes have relied on associations with genetic variants; this capacity to link genotype to phenotype has recently been accelerated by the plummeting cost of sequencing. The more recent advent of technologies which permit the quantification of endophenotypes like mRNA, metabolite, or protein abundance now enable mapping and trait dissection to be done between intermediate levels of biological organization. Assaying and associating these endophenotypes with traits of interest provides insight on biological mechanisms, serves as an independent source of evidence of associations, and facilitates prioritizing potentially causal variation while linking genes directly to traits in a way that potentially integrates the effects of multiple independent genetic variants. Here, we illustrated the utility of using a large RNA-seq resource in maize [17] for transcriptome-wide association studies and integrating these results with associations based on genetic variation.

We find evidence supporting the inclusion of transcriptome-wide variation in addition to genetic variation in models seeking to associate traits to underlying and likely causal genes in diverse maize lines, especially when the goal is to infer function of genes underlying oligogenic traits. Across tocochromanol trait classes, the inclusion of TWAS results enables more frequent detection of known causal genes and helps prioritize novel candidate genes in the profiled panel. Crucially, transcriptional variation alone does not improve over genotype-based associations, but it is in combination with genotypic information that the power of gene detection is increased.

As we demonstrate here, TWAS in combination with GWAS enhances the capacity to prioritize candidate genes over the use of GWAS alone. Given that more than half of detections are supported by TWAS (Table 1), our results also reveal much of the functional variation for these traits to be regulatory. While not all previously identified genes are detected by TWAS, this is likely a combination of insufficient power compared to the previous association studies in the NAM population with >16x as many observations [36], the sampling of a single time point per tissue, and the fact that not all functional variation is regulatory. Despite these limitations, TWAS adds value to GWAS mapping alone and increases the power to re-detect known genes. Our finding that TWAS alone is a valid method for finding true gene-trait associations is consistent with the recent findings of Lin and colleagues despite the difference between the eRD-GWAS and TWAS models [30]. However, our results differ in that we demonstrate that a combined test integrating TWAS and GWAS yields a more powerful test than either method individually when it comes to re-identifying known genes underlying oligogenic traits [36].

We also note that our efforts to validate our TWAS and GWAS detections differ from those of the previous study. In contrast to comparing the overlap of the detections by GWAS and TWAS in the same study, we compared our detections to previously known genes found in an independent set of germplasm, namely the NAM population [31], which was used to find tocochromanol associations [36]. Also, in contrast to the previously published study, we did not perform our cross-validation analysis in the same set of germplasm in which discovery was conducted by GWAS and TWAS to assess accuracy. Using variance partitioning in the independent NAM population, we found similar levels of variance explained by the genes detected by each method in the Goodman diversity panel [15], illustrating that even when the identified genes are tested in an outside population, the detections of the transcriptome-only and combined methods are found to be valid and explain similar amounts of variance to the genotype-based methods (Figs. 4, 5). This is roughly consistent with the cross-validation results comparing SNP_BayesB and eRD-GWAS presented in Table S4 by Lin and colleagues [30]. However, the previously published results show an advantage for eRD-GWAS for only one of fourteen traits, while on the basis of variance partitioning for kernel traits we find an advantage for the kernel based TWAS or the Fisher’s combined model for nine of the ten kernel-based traits for which measurements in NAM exist.

In further contrast to the previously published work [30], none of the SNPs used in our GWAS or variance partitioning methods were derived from RNA-seq data, allowing for a less bias towards expressed genes and giving the genotype-based tests more independence from the expression-based tests. In the previous work, more than 0.9M of the 1.2 M genetic variants were derived from the alignment of RNA-seq reads [30,37], potentially confounding the ability to make associations by GWAS with the presence of an expressed gene, and thus limiting the power of the genotype-based GWAS to make associations which are independent of expression.

It is striking that even in diverse maize lines where linkage decays quickly and thus the power to resolve mapping peaks to individual genes is high, TWAS provides a valuable supplement to genetic mapping alone. This benefit of TWAS would be compounded in species or populations in which resolution is limited. Additionally, by imputing expression values based on local/*cis* haplotype, as has been successfully shown in humans [22], the utility of TWAS could potentially be extended further in maize. Imputing expression to a larger panel would permit the exploitation of previously measured phenotypes across a much larger set of individuals which have not been expression profiled. By imputing only the local/*cis* genetic component of expression, and implicitly averaging over *trans* and environmental effects, the capacity to attribute field phenotypes to the genetic component of expression would likely be further improved.

The lack of improvement in re-detecting known tocochromanol traits by the multi-tissue TWAS models alone or as part of the Fisher’s combined tests is notable, but unsurprising for these genetically simple and very tissue specific traits. This lack of improvement indicates that kernel-based expression alone is most predictive of the kernel-based metabolites and accuracy is not improved by the incorporation of all other tissues. Rather than comparing the inclusion of all tissues vs kernels only, in the future a variable (tissue) selection TWAS approach should be used in which can remove uninformative terms from the model rather than including them but giving them a very small coefficient. It is also plausible that for more genetically complex traits which are also affected by expression across tissues, the multi-tissue TWAS results are more likely to be informative.

A further cause of the limited improvement for the kernel TWAS or Fisher’s combined test seen in the variance partitioning results is likely because GWAS identifies genomic regions which, when expanded to a 1 Mb window, could cover the functional variants. Furthermore, while the correct functional gene may not be prioritized by GWAS, if the trait is affected by genetic regulation rather than coding sequence change, the sites near the GWAS hit may in fact be more functional than those near the mechanistically significant gene itself even if they are misattributed to the incorrect proximal gene. Using a large independent diverse panel with very low LD to assess the heritability explained by the SNPs identified by each method may also provide a better estimate as the functional variants are not as easily tagged over long distances.

While the utility of expression endophenotypes in dissecting traits has been demonstrated here, it should be noted that associations made between endophenotypes and terminal phenotypes are inherently more susceptible to environmental effects than genotype-based associations. This susceptibility to environmental effects likely allows us to associate only the environmentally independent heritable fraction of expression with phenotype in our study, especially because expression data were collected from separate plants than those for which terminal phenotypes were measured. Given that in endophenotype-based association studies, like TWAS, environmental variation separately impacts and increases error in both the independent and dependent variables, methods like TWAS alone may plausibly be expected to perform more poorly than genetics-based associations. However, this shortcoming is partially compensated for by the more direct link between endophenotype and terminal phenotype and the potential discovery of mechanism. The collection of expression data from the same plants and conditions in which the phenotypes are collected would likely benefit the dissection of genotype by environment interactions by highlighting the impact of variation in expression for a specific gene within an environment, but cannot be examined here as terminal phenotypes and expression values were calculated from separate environments and years.

## Materials and Methods

### Genotypic data

Genotypes for the Goodman diversity panel [15] used in the genome-wide association studies were from the unimputed maize HMP 3.2.1 called against the B73 reference genome [38]. Variants segregating above 5% MAF in the union of all lines were considered for mapping. Variance component estimation in the maize NAM population [31] was performed using imputed HMP 3.2.1 variants [filename: NAM_HM321_KNN.hmp.txt.gz].

### Phenotypic data

For mapping in the Goodman diversity panel, kernel carotenoid BLUPs from 30 traits were from [28] and the 20 kernel tocochromanol traits BLUPs were from [27] after additional outliers were removed. The 22 field-based agronomic trait BLUPs were those calculated by [32]. Phenotypes used in variance partitioning with the maize NAM population were from [36] for the tocochromanol traits. Agronomic trait BLUPs were previously calculated by [32].

### Expression data

Expression quantifications were those created from seven diverse tissues in maize by aligning 3’ mRNAseq reads against the AGPv3.29 maize genome as described by [17].

### Genome-wide association study

Genome-wide association tests were conducted in the maize Goodman diversity panel [15] using a mixed linear model as implemented in FastLMM [39] accounting for kinship and a naive general linear model fit using MatrixEQTL [40] as implemented in TASSEL [41].

### Transcriptome-wide association study

Transcriptome-wide association tests were conducted in the maize Goodman diversity panel [15] for genes that were expressed in at least half of individuals represented in a specific tissue. A linear model was fit individually for each phenotype*expressed gene combination in which the explanatory variable is the expression value of a gene across individuals. TWAS was attempted both without covariates and with five genetic principal coordinates (calculated from maize HMP3.2.1 used in [17] and 25 PEER hidden factors (calculated separately for each tissue) as calculated in [17]. Multi-tissue TWAS was also performed. First a model was fit once per trait using the 5 principal coordinates described above. This model was then compared by ANOVA to a model for each gene containing terms for each tissue and the principle coordinates. The p-value resulting from this ANOVA was used to determine whether the multi-tissue model is significantly better than the covariate-only model. This p-value was also used as the p-value in the second of the Fisher’s combined tests below.

### Fisher’s combined tests of TWAS and GWAS

The GWAS p-value (mixed linear model with kinship as a random effect) of each SNP in the top 10% of most associated SNPs was assigned to nearest gene and then combined with the TWAS p-value (linear model with MDS PCs + PEERs) for that same gene using Fisher’s combined test as implemented in the sumlog method in the *metap* package [42] in R. TWAS p-values for genes which were not tested in TWAS (i.e. their expression was not observed in at least half of individuals) were set to p=1 prior to combining with GWAS p-values. Fisher’s combined tests were performed in the same way when including the multi-tissue TWAS results instead of the kernel-only results.

### Variance partitioning

Using the KNN imputed Nested Association Mapping population HMP3.2.1 genotypes described above, kinship matrices were calculated based on the top ten genes identified by each of the TWAS, GWAS, and combined models described in the Goodman diversity panel [15]. To independently assess the accuracy of detected genes, the phenotypic variance explained by each kinship matrix was calculated in the Nested Association Mapping population, within each family and across all the NAM families. For TWAS, the top 10 genes were taken and all SNPs within a 0.5 Mb radius of the start and end of the gene (maize annotation AGPv3.29) were used to calculate a single kinship matrix per trait using the Variance Component Annotation Pipeline in TASSEL [41]. The REML solver in LDAK [35] was used to calculate the variance explained by the single kinship matrix. For GWAS, the SNPs were ordered based on significance and assigned to their nearest gene. The top ten unique genes from this list were taken to calculate kinship matrices using the same 0.5 Mb radius around the gene. To avoid picking multiple genes and redundant variants from the same peak based the GWAS results, the top most associated gene was used within a peak and all other genes within the 0.5 Mb radius were excluded from selection as top genes.

### Overlap with known kernel metabolite genes

Fourteen known tocochromanol biosynthetic genes identified in NAM [36] and 58 *a priori* candidate genes relevant to the biosynthesis and retention of carotenoids [28] were used as positive controls to test the capacity of our GWAS, TWAS, and combined methods to re-detect known genes. In order to avoid comparison of p-value thresholds across methods, positive detections were counted if a gene was detected among the top 1% of genes associated with a trait.

## Author contributions

KAGK performed transcriptome-wide associations (TWAS), joint tests of TWAS and genome-wide associations (GWAS) results, and quantified overlaps with known genes. NBB performed GWAS mapping and variance partitioning. CHD and MAG developed and provided lists of identified tocochromanol and carotenoid associated genes from NAM and developed a filtered set of carotenoid and tocochromanol phenotypes for NAM and the Goodman diversity panel. KAGK, NBB, and ESB conceived of the study. KAGK and NBB wrote the manuscript.

## Acknowledgements

We thank Sara Miller for thorough reading of the manuscript and assistance with syntax. This work was supported by the US Department of Agriculture–Agricultural Research Service and the National Science Foundation grants IOS-0922493 and IOS-1238014 to E.S.B. The National Science Foundation Graduate Research Fellowship Program grant DGE-1650441 and the Section of Plant Breeding and Genetics at Cornell University provided support to K.A.G.K.

---

**Figure S1.**
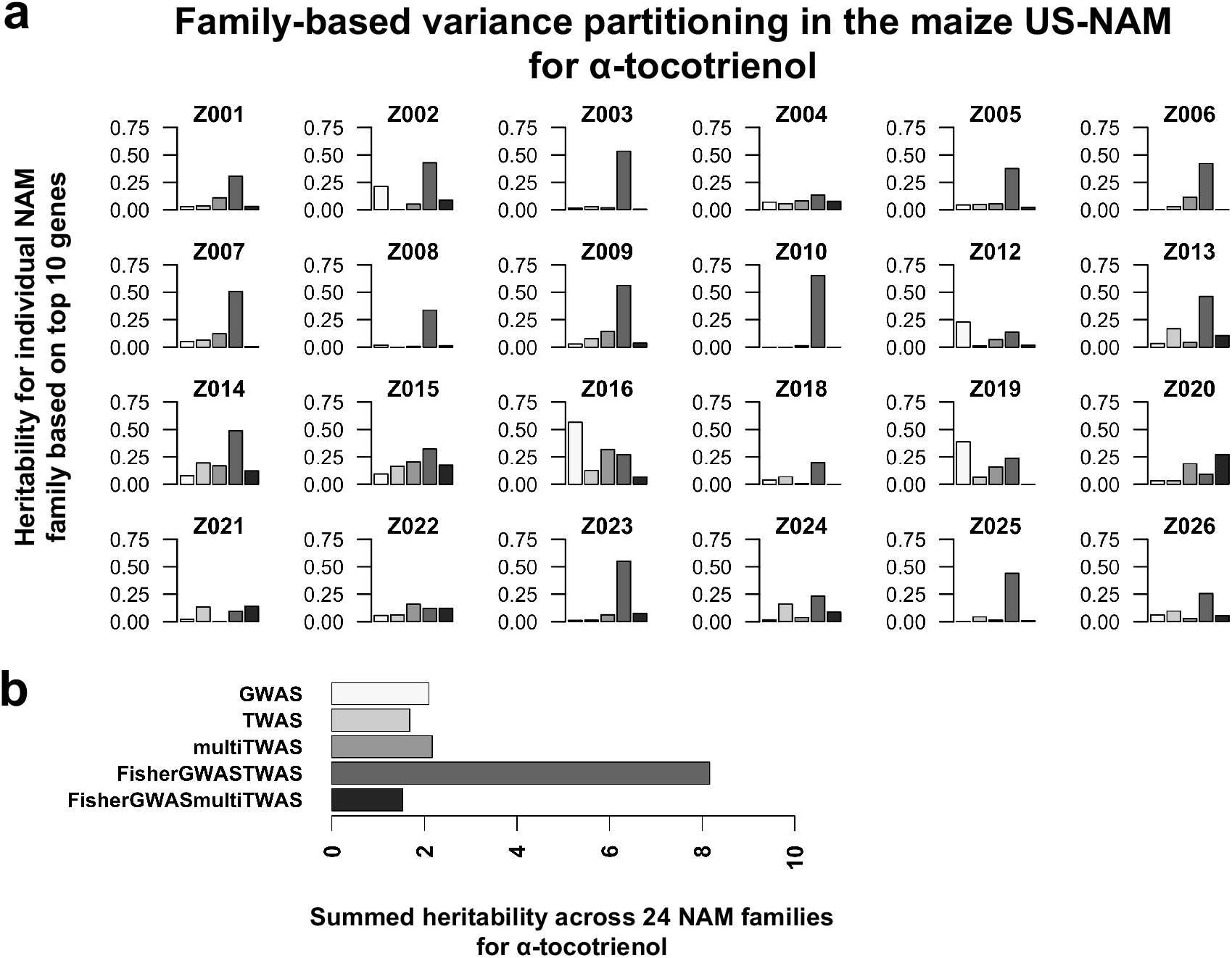
Family-based variance partitioning on individual NAM family for alpha-tocotrienol. This is an example in which the Fisher’s combined test of the MLM GWAS + kernel-based TWAS explained higher heritability relative to other models. **a** Heritability was estimated for each of 24 NAM families using kinship matrices made from the genetic regions adjacent to the top 10 ranked genes mapped by MLM GWAS, kernel-based TWAS, multi-tissue TWAS, the Fisher’s combined test of the MLM GWAS + kernel-based TWAS, and the Fisher’s combined test of the MLM GWAS + multi-tissue TWAS. **b** Heritability estimates were then added together (hereafter, summed heritability). Different models were compared in horizontal barplot based on summed heritability explained in 24 NAM families.

**Figure S2.**
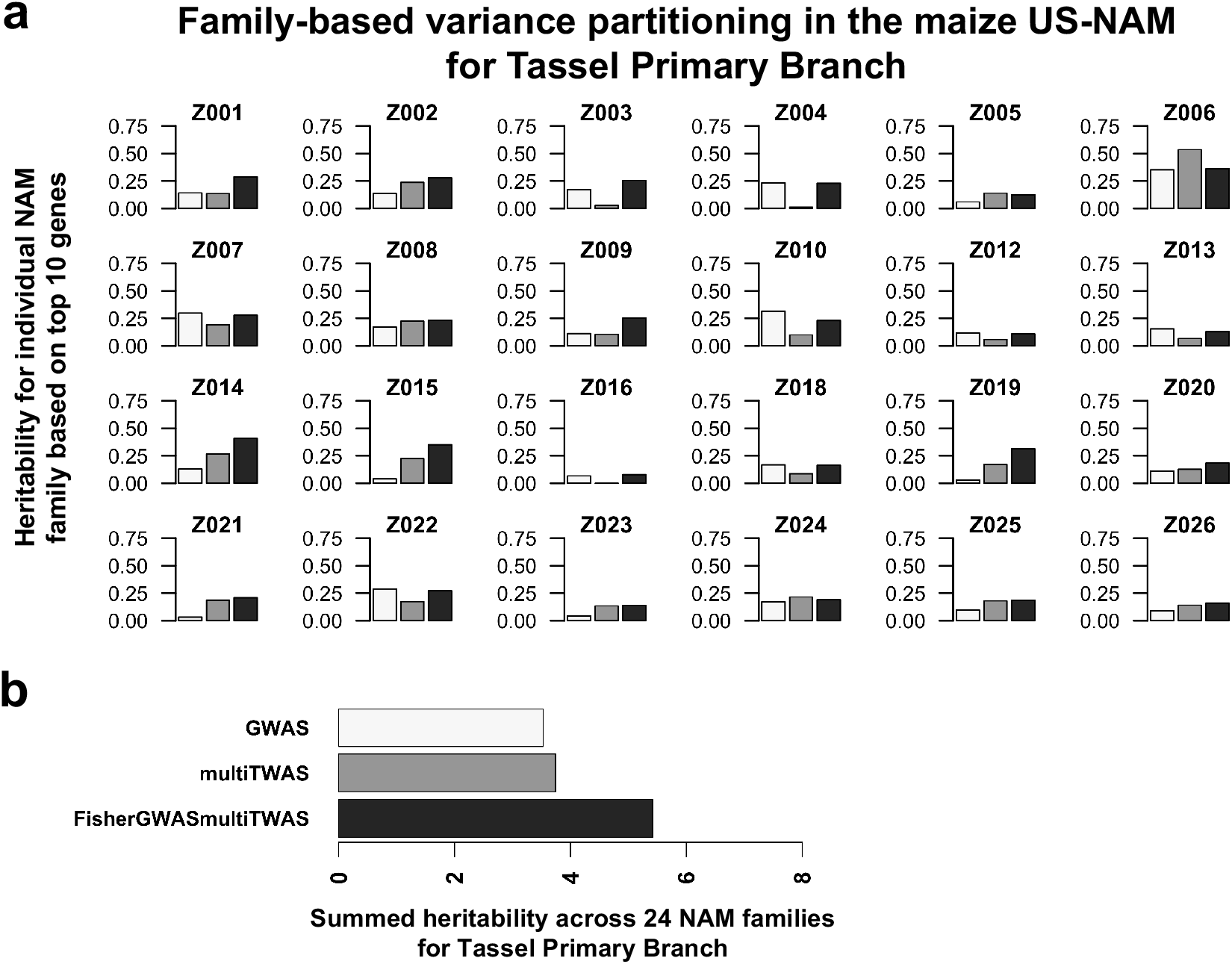
Family-based variance partitioning on individual NAM family for tassel primary branch. This is an example in which the Fisher’s combined test of the MLM GWAS + multi-tissue TWAS explained higher heritability relative to other models. **a** Heritability was estimated for each of 24 NAM families using kinship matrices made from the genetic regions adjacent to the top 10 ranked genes mapped by MLM GWAS, multi-tissue TWAS, and the Fisher’s combined test of the MLM GWAS + multi-tissue TWAS. **b** Heritability estimates were then added together (hereafter, summed heritability). Different models were compared in horizontal barplot based on summed heritability explained in 24 NAM families.

**Figure S3.**
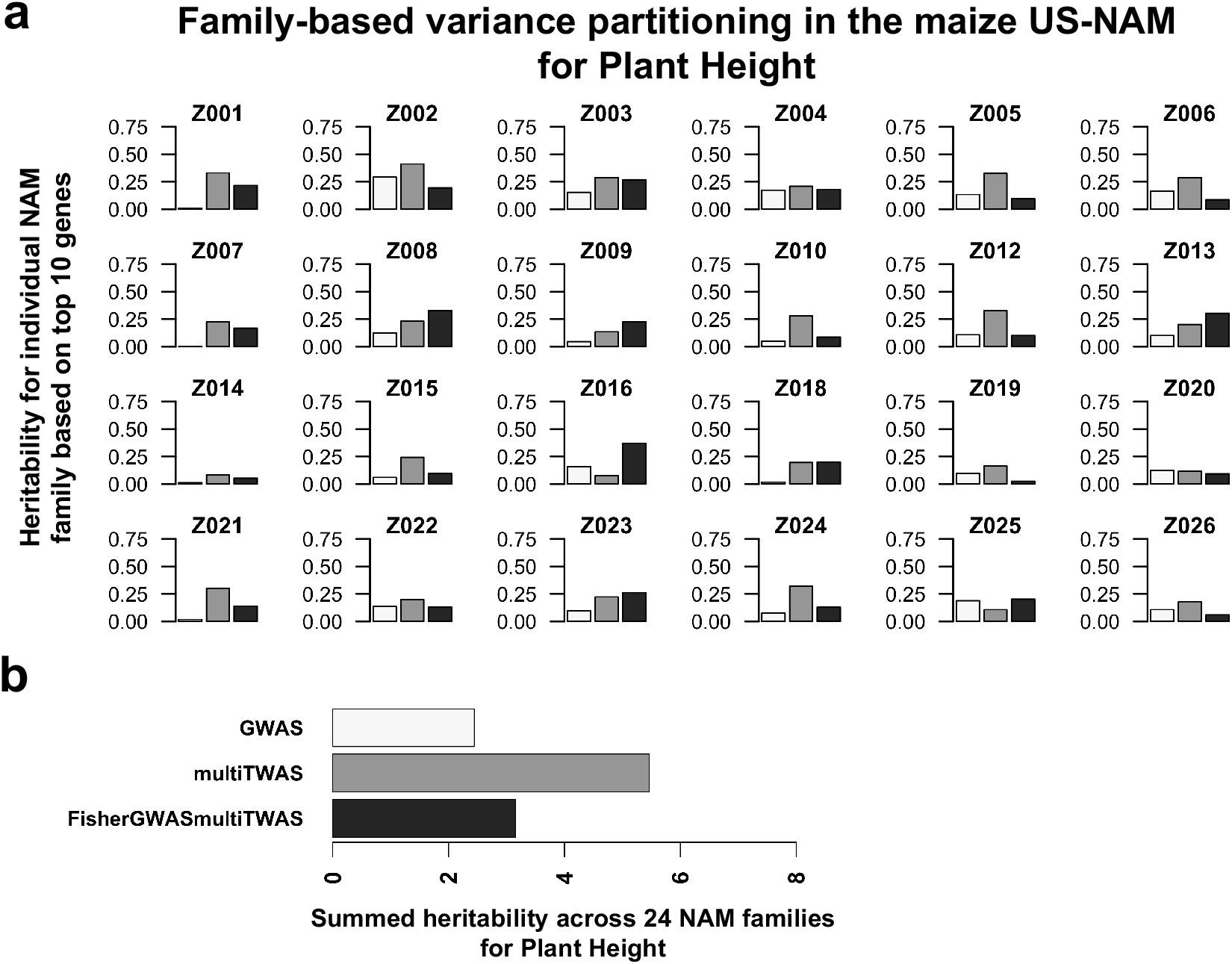
Family-based variance partitioning on individual NAM family for plant height. This is an example in which the multi-tissue TWAS explained higher heritability relative to other models. **a** Heritability was estimated for each of 24 NAM families using kinship matrices made from the genetic regions adjacent to the top 10 ranked genes mapped by MLM GWAS, multi-tissue TWAS, and the Fisher’s combined test of the MLM GWAS + multi-tissue TWAS. **b** Heritability estimates were then added together (hereafter, summed heritability). Different models were compared in horizontal barplot based on summed heritability explained in 24 NAM families.

**Table S1.** Percentile ranks from kernel-based TWAS, multi-tissue TWAS, MLM GWAS, Fisher’s combined (kernel-based TWAS + MLM GWAS), and Fisher’s combined (multi-tissue TWAS + MLM GWAS) results for genes previously identified in NAM for tocochromanol traits in maize [36].

**Table S2.** Percentile ranks from kernel-based TWAS, multi-tissue TWAS, MLM GWAS, Fisher’s combined (kernel-based TWAS + MLM GWAS), and Fisher’s combined (multi-tissue TWAS + MLM GWAS) results for carotenoid candidate gene list in maize as described by [28].

